# Ultrasonic measures of prestin (SLC26a5) charge movements in membrane patches

**DOI:** 10.1101/2022.10.03.510610

**Authors:** Joseph Santos-Sacchi, Jun-Ping Bai, Dhasakumar Navaratnam

## Abstract

Charged moieties in the outer hair cell (OHC) molecular motor protein, prestin, are driven by transmembrane voltage to ultimately provide for cochlear amplification. The speed of voltage-dependent conformational switching underlies its ability to influence micromechanics of the cell and the organ of Corti. Corresponding voltage-sensor charge movements in prestin, classically assessed as a voltage-dependent, nonlinear membrane capacitance (**NLC**), have been used to gauge its frequency response. Using megahertz sampling of prestin charge movements, we extend interrogations of prestin performance into the ultrasonic range (up to 120 kHz) and find response magnitude larger than previously reported. We also confirm kinetic model predictions of prestin by directly observing its cut-off frequency under voltage-clamp as the intersection frequency (F_is_) of the real and imaginary components of complex **NLC (cNLC)**, showing values near 19 kHz. At higher frequencies, the imaginary component roll-off exactly tracks that of Abs(**cNLC**). The frequency response of prestin displacement current noise determined from the Nyquist relation aligns with this cut-off. On the other hand, previous measures of stationary thermal-driven noise of prestin indicated that the cut-off was several fold greater than that of **NLC**, in violation of the fluctuation-dissipation theorem. We have attempted to confirm this apparent paradox, but find that low frequency (<10kHz), voltage-dependent 1/f noise, likely due to intrinsic prestin conductance, can limit the accessible bandwidth for stationary noise analysis. Nevertheless, within those bandwidths, frequency response comparisons of stationary measures and Nyquist relation measures are consistent. We conclude that voltage stimulation accurately assesses the spectral limits of prestin activity.

**Significance:** Using megahertz sampling, we extend measures of prestin charge movement into the ultrasonic range and find that the frequency roll-off is less than previously reported. Nevertheless, analysis of complex nonlinear capacitance confirms low-pass behavior, with a characteristic cut-off frequency near 19 kHz. The frequency response of prestin noise garnered by the admittance-based Nyquist relation confirms this cut-off frequency. In conflict with previous results, however, we find a similar low-pass frequency response using direct measures of prestin noise in the absence of voltage stimulation. Our data indicate that voltage perturbation provides an accurate assessment of prestin performance.

## Introduction

Voltage is the physiological driving force that underlies prestin activity (Santos-Sacchi and Dilger, 1988) and mammalian cochlear amplification. That is, AC receptor potentials of outer hair cells (OHC) evoked by stereociliary bundle displacements across the acoustic spectrum will drive prestin’s conformational switching and thereby alter OHC electro-mechanics leading to enhanced hearing (cochlear amplification) (Dallos et al., 2008). Voltage-clamp studies on the electro-mechanical activity of the isolated OHC and patches of its lateral membrane have led the way in defining prestin’s frequency performance by measuring electromotility directly and/or correspondingly the protein’s voltage-sensor activity (nonlinear capacitance, **NLC**). At the voltage where prestin sensor charge is equally distributed across the membrane, namely V_h_, the midpoint of its Boltzmann distribution, a characteristic kinetic response is obtained whose cut-off frequency is equal to twice the forward or backward transition rate, the rates being equal at this voltage. Each measure of prestin performance at V_h_ has been shown in recent years to be low pass in nature, rolling off significantly above 10-20 kHz (see (Santos-Sacchi and Tan, 2022)). On the other hand, auditory perception in some mammals extends into the ultrasonic range, even beyond 100 kHz.

Prestin has dual sensitivity to voltage and membrane tension (Iwasa, 1993; Gale and Ashmore, 1994; Kakehata and Santos-Sacchi, 1995; Ludwig et al., 2001; Santos-Sacchi et al., 2001), like some ion channels (Beyder et al., 2010; Schmidt et al., 2012; Jin et al., 2020; Perez-Flores et al., 2020). Whereas prestin has been uniquely described as a piezoelectric device, those other dually sensitive proteins have not. Consequently, concern has been raised that the measured kinetics (frequency response) of prestin suffers from untoward mechanical influences of measurement techniques; notably, for macro-patch experiments the pipette fluid column load has been suggested to slow down prestin kinetics (Dong et al., 2000). However, these concerns have been directly refuted, and instead it was shown that the frequency response of prestin’s **NLC** using this technique registers alterations induced by such physiologically important factors as intracellular chloride level and membrane fluidity/viscosity, even revealing interactions between membrane viscosity and membrane tension (Santos-Sacchi and Tan, 2022). Thus, the kinetics of prestin, as expected with any membrane-bound, voltage-dependent protein, depends on the influence of its local molecular environment. Indeed, prestin-prestin interactions themselves, occurring when prestin membrane density is sufficiently high, likely account for negative cooperativity that influences the distribution of prestin conformations (Zhai et al., 2020), and possibly sets the foundation for mechano-sensitivity in prestin, as opposed to piezoelectricity, per se. Indeed, the sub-nanometer cryo-EM structure of prestin does not indicate any major difference between the non-electromechanical SLC26 family member SLC26a9 and prestin (RSMD ~3Å) that could be taken as evidence for an actual piezoelectric device (Bavi et al., 2021; Ge et al., 2021; Butan et al., 2022).

To reiterate, the speed limit of prestin’s conformational switching cannot be divorced from the influence of its native environment, and the physiologically important limit must be that resulting from the natural stimulus, namely voltage. The seminal observations of Gale and Ashmore (Gale and Ashmore, 1997) using voltage-clamped membrane patches characterized **NLC**, determined at V_h_, as low-pass. Surprisingly, subsequent measures of electromotility (**eM**) contrarily suggested that responses driven by voltage are ultrasonic (Frank et al., 1999). These **eM** measures were made at resting potentials far removed from V_h_. In fact, **eM** has since been shown to be low pass at V_h_, exactly where the characteristic low-pass **NLC** frequency response is appropriately measured (Santos-Sacchi and Tan, 2018). Nevertheless, it has been suggested that the kinetics of thermal noise of prestin in membrane patches defines the protein’s limiting speed, and that the frequency cut-off is much higher than that of voltage-driven, admittance-based measures of **NLC** in those same membrane patches (Dong et al., 2000). That work indicated a violation of the fluctuation-dissipation theorem, which posits that the frequency response of voltage-driven, admittance-based noise estimates and those made in the absence of voltage stimulation (thermally driven at equilibrium) are the same (Nyquist, 1928; Callen and Greene, 1952; Lauger, 1978). The violation was rationalized by speculating that mechanical influences on prestin movements are different in the absence of voltage stimulation. Included in that rationalization was the suggestion that patch pipette column load lowers prestin’s frequency response, which as noted above is not the case (Santos-Sacchi and Tan, 2022). So, why do stationary thermal-driven noise measures differ from estimates based on the admittance-based Nyquist relation? Here we reinvestigate this seeming paradox, while providing new ultrasonic measures of prestin charge movement that differ from previous reports.

## Materials and Methods

OHC complex **NLC** (**cNLC**) was measured in macro-patches under voltage clamp as previously described (Santos-Sacchi and Tan, 2020; Santos-Sacchi et al., 2021), except rather than sample currents at 100 kHz, we now sampled currents at 1 MHz (16 bit NI-USB 6356; National Instruments) using an Axon 200B amplifier (capacitive feedback mode) with Bessel filter set at 100 kHz. Briefly, macro-patches were established on the OHC lateral plasma membrane, and stimulus partitions with and without AC voltage chirps (10 mV) were presented atop holding potentials ranging from +160 to −160 mV (see **Fig. 1**). Currents were corrected for system roll-off characteristics in the frequency domain. Real and imaginary components of **cNLC** were extracted following the methodology of Fernandez et al. (Fernandez et al., 1982). In general, complex capacitance (cC_m_) is derived from membrane admittance, *Y*_*m*_(*ω*)

**Fig. 1.**
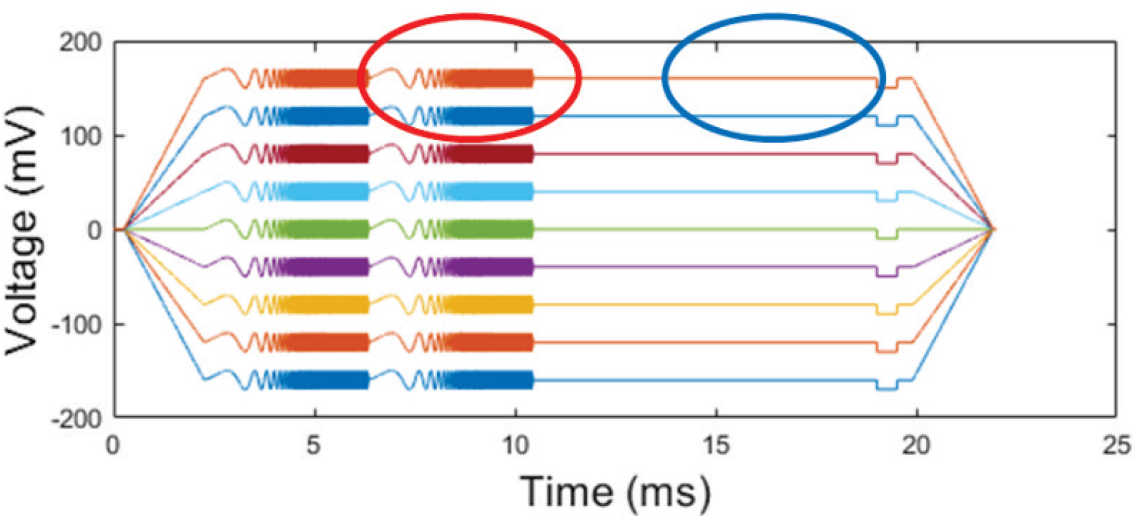
Voltage-clamp protocol to measure **NLC** and admittance-based noise estimates (Nyquist relation, *eq. 1*) using voltage chirp stimuli (red circle highlight), and thermally driven noise (*eq. 2*) in the absence of voltage perturbation (blue circle highlight). Holding voltage was ramped up from +160 mV to −160 mV in 40 mV increments. The protocol was repeated 64x and individual spectra averaged real time in jClamp.

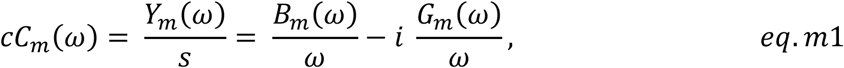

providing real and imaginary components,

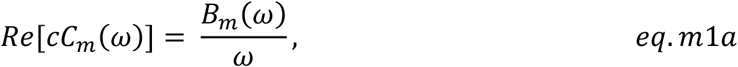

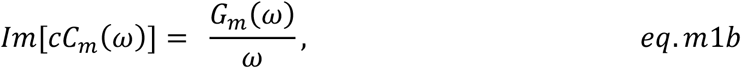

where B_m_ is membrane suseptance, G_m_ is membrane conductance, *s=iω, ωω* = 2π and 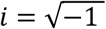 The complex components represent voltage sensor charge moving in and out of phase with AC voltage, and Im(cC_m_) does *not* represent an actual membrane conductance. **cNLC** and admittance-based (Nyquist relation) prestin noise estimates were obtained following subtraction of linear admittance at +160 mV holding potential, where OHC **NLC** is absent. Prestin noise spectra were directly measured at those holding potential partitions lacking AC voltage stimulation (see **Fig. 1**). In this case, the spectrum at +120 mV was subtracted from all others. Spectra were averaged 64 times, using 4096 data points.

We fit **NLC** as a molecular capacitor obeying Boltzmann statistics (Santos-Sacchi, 1991; Santos-Sacchi and Navarrete, 2002).

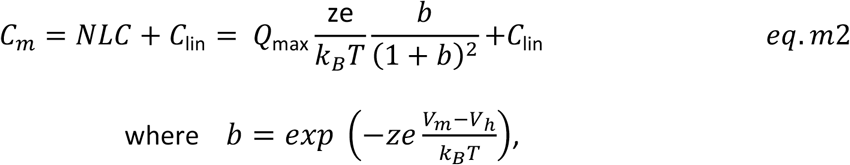

Q_max_ is the maximum nonlinear charge moved, V_h_ is voltage at peak capacitance or equivalently, at half-maximum charge transfer, V_m_ is membrane potential, *z* is valence, C_lin_ is linear membrane capacitance, e is electron charge, *k*_*B*_ is Boltzmann’s constant, and T is absolute temperature. Fits were made to real and imaginary components of **cNLC** across interrogation frequency.

Analyses were made in jClamp (www.scisoftco.com) and Matlab (www.mathworks.com). Methods have been approved by Yale’s animal use committee.

## Results and Discussion

The fluctuation-dissipation theorem indicates that the frequency response of admittance-based noise estimates (Nyquist relation; *eq*.*1*) and those made in the absence of voltage stimulation (stationary, thermally-driven at equilibrium; *eq. 2*) are the same (Nyquist, 1928; Callen and Greene, 1952).

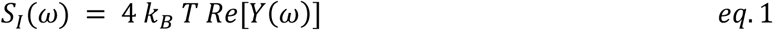

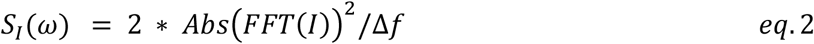

where *Y* and *I* are admittance and current, respectively. *⊿f* is number of sampled points, *npts* (4096 in our experiments) times sampling rate, *F*_*s*_ (1 MHz in our experiments); ⊿*f* = *npts* * *F*_*s*_. The other symbols have their usual meaning. In order to measure admittance-based and stationary noise concurrently, we utilized the protocol shown in **Fig. 1. NLC** and Nyquist noise spectra were simultaneously assessed with voltage chirps; thermal noise was assessed in the absence of AC voltage perturbation, all within the same voltage-clamp protocol.

The violation of the fluctuation-dissipation theorem is sometimes observed biophysically, and may indicate either that noise measures were not made at equilibrium, or that modes of elementary motions are different depending on internal (thermal) or external (voltage) excitations (Fishman, 1985). Dong et al. have argued the latter to account for their disparate observations on prestin voltage-sensor kinetics for **NLC** and thermally-driven fluctuations (Dong et al., 2000).

Kolb and Lauger (Kolb and Lauger, 1977) in their derivation of voltage-sensor noise in a multi-state (or 2-state) model, showed formal equivalence to a series RC model, each showing an inverse Lorentzian response. We illustrate the two methods of noise assessment, admittance-based Nyquist relation (*eq*.*1*) and direct stationary noise measurement (*eq*.*2*), in **Fig. 2**, using a physical series RC model.

**Fig. 2.**
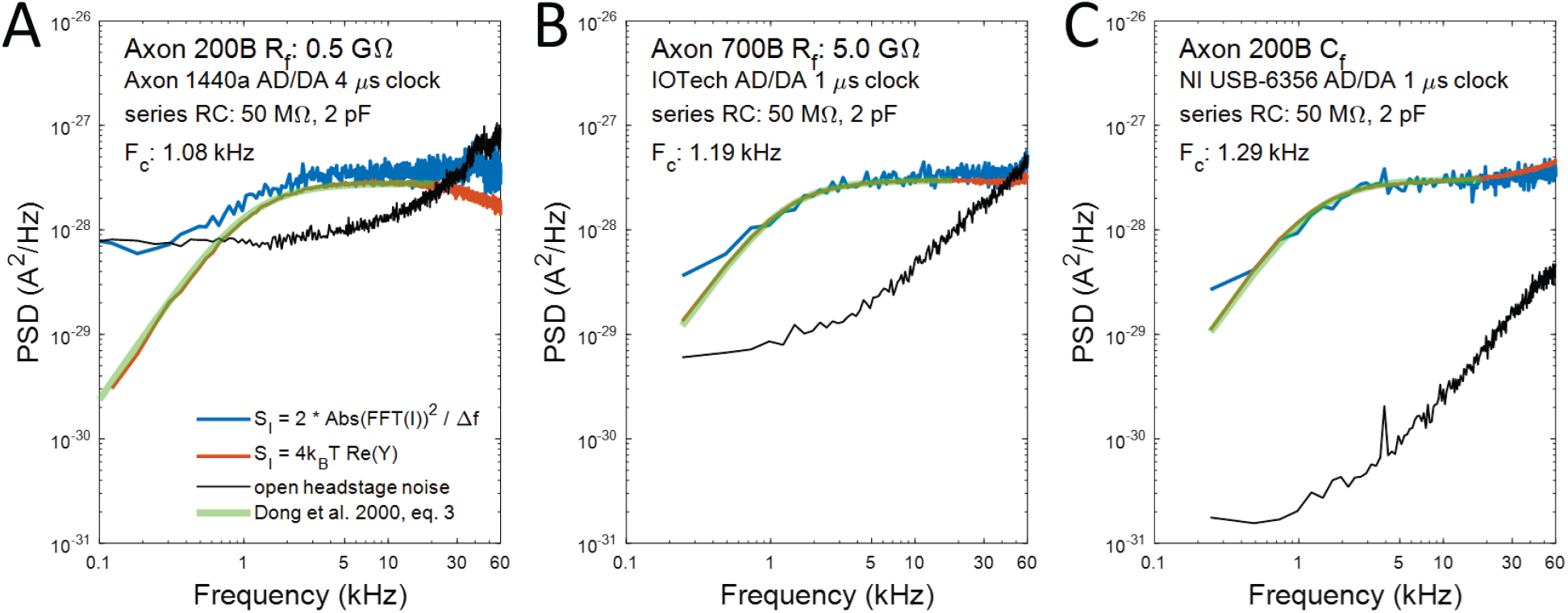
Noise in series RC circuit illustrating measurements made with (admittance-based, red line) or without (direct, blue line) voltage perturbation, employing **A)** Axon 200B amplifier with 0.5 GΩ feedback resister, **B)** Axon 700B amplifier with 5.0 GΩ feedback resister, and **C)** Axon 200B amplifier with feedback capacitor. Fits were made to the admittance-based measures using *eq. 3* from Dong et al. (2000) (green lines). Cut-off frequency for each method is essentially the same. The feedback component determines the headstage noise (black lines), with capacitive feedback far superior to resistive feedback. Our measure of 200B headstage noise with 0.5 GΩ feedback is equivalent to that published by Rosenstein et al. (Rosenstein et al., 2012). We use capacitive feedback to avoid confounding headstage noise.

**Fig. 2** shows, using two different patch clamp amplifiers (Axon 200B and 700B), that the two methods (*eq*.*1* or *eq. 2*) to measure noise in a series RC circuit are equivalent. The frequency response is an inverse Lorentzian, providing essentially the same cut-off frequency for either method, where the response magnitude reaches 0.707 that of the high frequency asymptote. The fitted inverse Lorentzian curves (green lines) were obtained with previously derived equations (Kolb and Lauger’s *eqs. 8-9* (Kolb and Lauger, 1977); or equivalently, Dong et al.’s *eq. 3*. (Dong et al., 2000), to provide the cut-offs (F_c_) listed in the three plots. The 200B amplifier feedback resistor (R_f_) of 0.5 GΩ is the same setting used by Dong et al. (Dong et al., 2000). The 700B amplifier feedback resistor (R_f_) was set to 5 GΩ· Additionally, the 200B was used in capacitive feedback mode. Baseline headstage noise (black lines), given by 4K_B_T/R_f_, differs for these three setups (**Fig.2 A, B, C**), with capacitive feedback far superior to resistive feedback. Amplifier parasitic capacitance contributes to the high frequency noise evident in all feedback modes, and is not responsive to amplifier front-panel capacitance compensation. Our measure of 200B headstage noise with 0.5 GΩ feedback (**Fig. 2 A**, black line) is equivalent to that published by Rosenstein et al. (Rosenstein et al., 2012). Data presented below utilize the capacitive feedback setup to avoid limiting headstage noise. As will be shown below, besides headstage and R_f_ noise, low frequency 1/f noise will greatly impact the ability to directly (*eq. 2*) analyze thermally-driven prestin noise levels. On the other hand, admittance-based noise estimates (*eq. 1*) are readily achievable because of enhanced signal-to-noise afforded by voltage stimulation.

Prestin **NLC** is measurably a complex quantity, **cNLC** (Santos-Sacchi et al., 2021), separable into real and imaginary components. In **Fig. 3**, these components are plotted as a function of holding voltage and frequency. Fits (examples in red) are made with *eq. m2* and provide an average V_h_ of −82.77 +/−7.62 mV @ 3.66 kHz (n=9). Below a particular frequency (F_is_, see below), there is a decrease in the positive real component accompanied by an increase in the negative imaginary component, most notable at V_h_. Here we have extended our bandwidth beyond our previous 30 kHz bandwidth (Santos-Sacchi and Tan, 2020), which now enables us to discern the peak and subsequent reduction in the negative imaginary component as frequency increases; this was not observable in our earlier lower bandwidth measures (Santos-Sacchi and Tan, 2020; Santos-Sacchi et al., 2021).

**Fig. 3.**
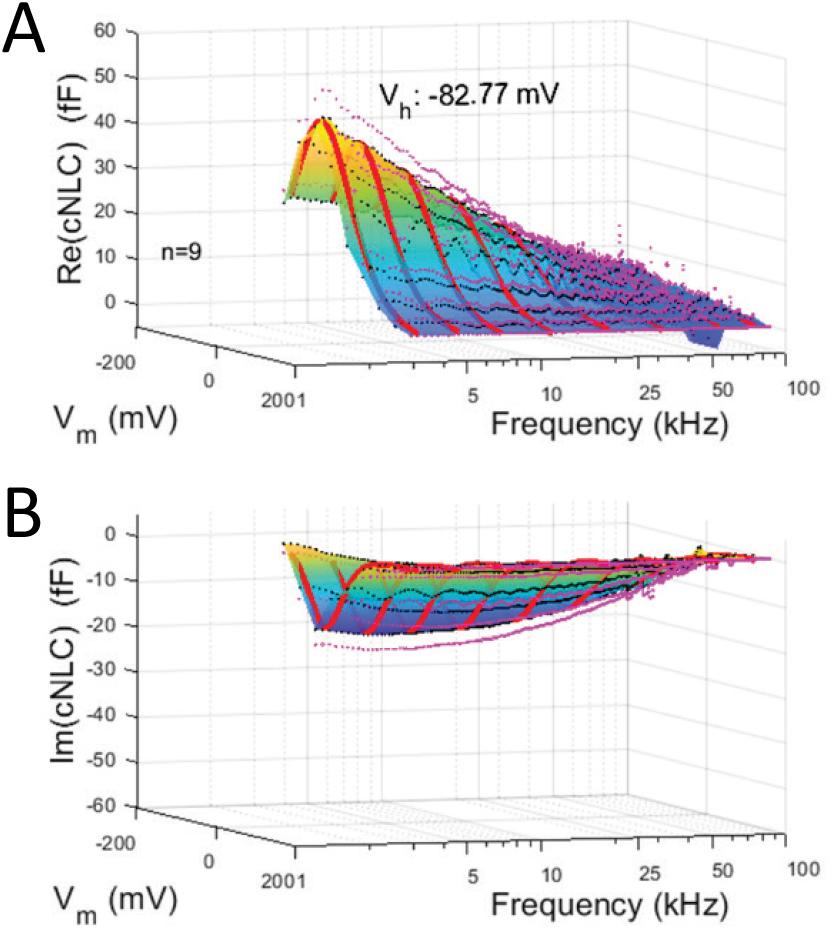
Real (**A**) and imaginary (**B**) components of OHC complex **NLC** displayed out to 100 kHz. Fits were made with the 2-state Boltzmann *eq. m2*, with black dots showing the means, and magenta dots showing the SEM of 9 averaged patches. Example fits are shown in red. Of notable importance at this recording bandwidth is the previously unobserved decrease in the imaginary component of **cNLC** above 30 kHz.

**Figs. 4 A, B, C** show the relationships of these complex components and Abs(**cNLC**) 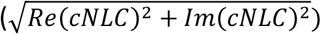 across frequency, from 5 to 120 kHz, at three different holding potentials, namely, −40 mV, −80 mV and −120 mV. The first and last holding potentials bracket the average V_h_ of – 82.77 mV. The imaginary component is multiplied by −1 to enable logarithmic plotting and comparisons. As can be seen, Abs(**cNLC**) shows the widest frequency response, well beyond the real component. The imaginary component peaks near a frequency (F_is_) where it intersects the real component. In a 2-state kinetic model, F_is_ corresponds to the −3dB point of Abs(**cNLC**) (see Appendix in Santos-Sacchi and Tan (Santos-Sacchi and Tan, 2022)). Importantly, at low frequencies Abs(**cNLC**) coincides with the real component, and at high frequencies Abs(**cNLC**) coincides with the imaginary component.

**Fig. 4.**
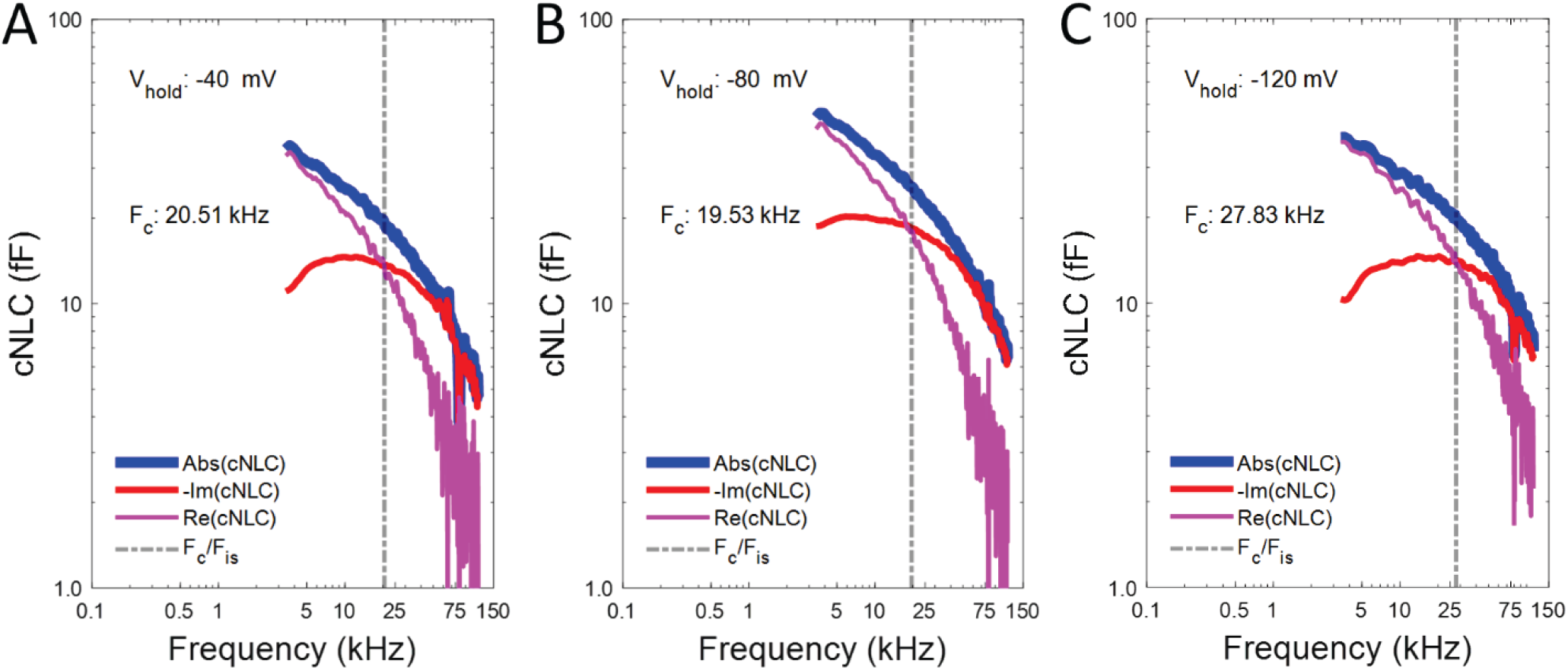
Components of **cNLC**, absolute, real and imaginary, as a function of stimulus frequency at holding potentials of A) −40 mV, **B)** −80 mV and **C)** −120 mV. The intersection frequency (F_is_) between real and imaginary components defines the characteristic roll-off of Abs(**cNLC**) across frequency. F_is_ increases as holding potential is displaced from V_h_, whose average for these cells is −82.09 mV. Indeed, Abs(**cNLC**) at −80 mV (**B**), essentially at V_h_, derived from real and imaginary components provide the lowest-pass response (blue solid line), which agrees well with previously published data. At this measurement bandwidth, it is revealed that Abs(**cNLC**) coincides with the imaginary component at high frequencies. Thus, the frequency response of Abs(**cNLC**) provides full details of the imaginary component’s influence at very high frequencies.

Thus, the frequency response of Abs(**cNLC**), which we have used to characterize prestin’s high frequency influence on cochlear mechanics (Santos-Sacchi and Tan, 2020; Santos-Sacchi et al., 2021), exactly characterizes the imaginary component at high frequencies. That is, if the imaginary component of **cNLC** is important for power production of prestin (Rabbitt, 2020; Rabbitt, 2022), its influence diminishes rapidly in line with Abs(**cNLC**) at high frequencies.

**Fig. 5** plots Abs(**cNLC**) near V_h_, (thick blue solid line) along with previous data from Gale and Ashmore (Gale and Ashmore, 1997) (magenta circles) and Santos-Sacchi and Tan (Santos-Sacchi and Tan, 2020) (green asterisks). The frequency response of our current dataset corresponds well to that of our scaled previous dataset (green asterisks) but extends our analysis of Abs(**cNLC)** out to 120 kHz. Additionally, the unified power fit from Santos-Sacchi and Tan (Santos-Sacchi and Tan, 2020) is plotted (dashed magenta line). That power fit was made to a combined data set of ours and Gale and Ashmore’s, which included the very high frequency responses (> 25 kHz) that they originally noted as adversely affected by amplifier limitations. We had argued that those high frequency responses were valid and thus used them in our unified fit. However, now that we have extended our bandwidth, we find that prestin charge movement does not roll off as precipitously as the power fit indicated, instead showing larger responses in the ultrasonic range.

**Fig. 5.**
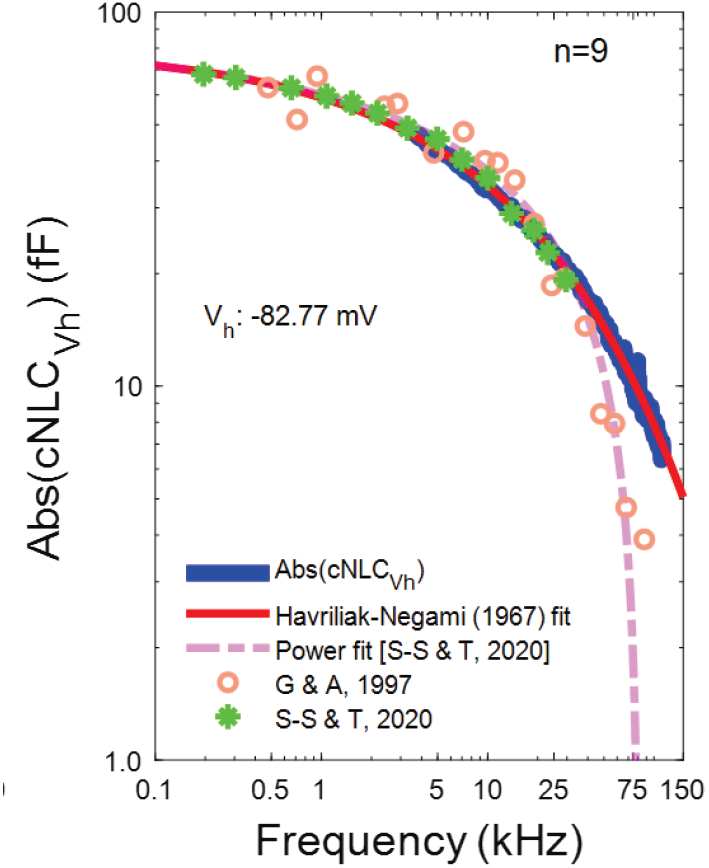
Prestin charge moves at ultrasonic frequencies. Near V_h_ (−80 mV holding potential), Abs(**cNLC**) defines the characteristic speed of prestin. Extension of assessment out to 120 kHz redefines its high frequency response, where it is necessary to fit the response not with a power function of frequency, but with the Havriliak-Negami [HN] relaxation function, the frequency domain equivalent of a stretched exponential response.

We now use a fit with the Havriliak-Negami [HN] relaxation function (Alverez et al., 1991), the frequency domain equivalent of a stretched exponential (red line). We have previously reported on stretched exponential relaxations of prestin’s **NLC** in response to voltage steps (Santos-Sacchi et al., 1998;

Santos-Sacchi et al., 2009; Santos-Sacchi and Song, 2014a). Indeed, our *meno presto* kinetic model incorporates this stretched exponential behavior and accurately describes a variety of prestin’s biophysical behaviors (Santos-Sacchi and Song, 2014b). Thus, the HN and power fits differ above 80 kHz, with the HN fit extending to higher frequencies. As we noted previously, either fit is indistinguishable for whole cell **NLC** measures out to 15 kHz (Bai et al., 2019). Based on our new wider bandwidth measures, there is a 20.5 dB drop at 120 kHz, or a 10.6 fold reduction in total (both in and out of phase with voltage) sensor charge movement. Nevertheless, F_is_ (or equivalently, F_c_) accurately describes the characteristic low-pass cutoff of Abs(**cNLC**), which is slowest at V_h_, and increases in cut-off frequency at holding voltages offset from V_h_. This type of frequency-dependent behavior relative to V_h_ is also observed in voltage-driven **eM** (Santos-Sacchi et al., 2019), where the lowest-pass **eM** response is likewise at V_h_.

Similar to the series RC noise measures above (**Fig. 2**), these admittance-based data can be used to estimate prestin thermal noise based on the Nyquist relation (*eq. 1*), and the frequency cut-off based on these noise measures should correspond to that of **NLC** F_c_ or F_is_. **Fig. 6** shows noise estimates and fits with *eq. 3* from Dong et al. (2000) of the averaged responses (solid gray lines), indicating that fitted noise cut-offs closely correspond to those of **NLC**, e.g., at −80 mV **NLC** F_c_ = 19.53 kHz and noise F_c_= 19.71 kHz; at −120 mV **NLC** F_c_ = 27.83 kHz and noise F_c_= 25.88 kHz. Mean and SEM of fits from individual macro-patches (n=9) are −80 mV: 20.01 +/−0.93 kHz; −120 mV: 26.49 +/−1.38 kHz; and −160 mV: 39.31 +/−3.43 kHz. Note that fits deviate from data at low frequencies likely because prestin behavior presents not exactly as a 2-state process, but more like a stretched exponential process (see above). Thus, prestin frequency responses based on either **NLC** or noise estimates using the Nyquist relation correspond, and at resting potentials away from V_h_ will be significantly faster than at V_h_.

**Fig. 6.**
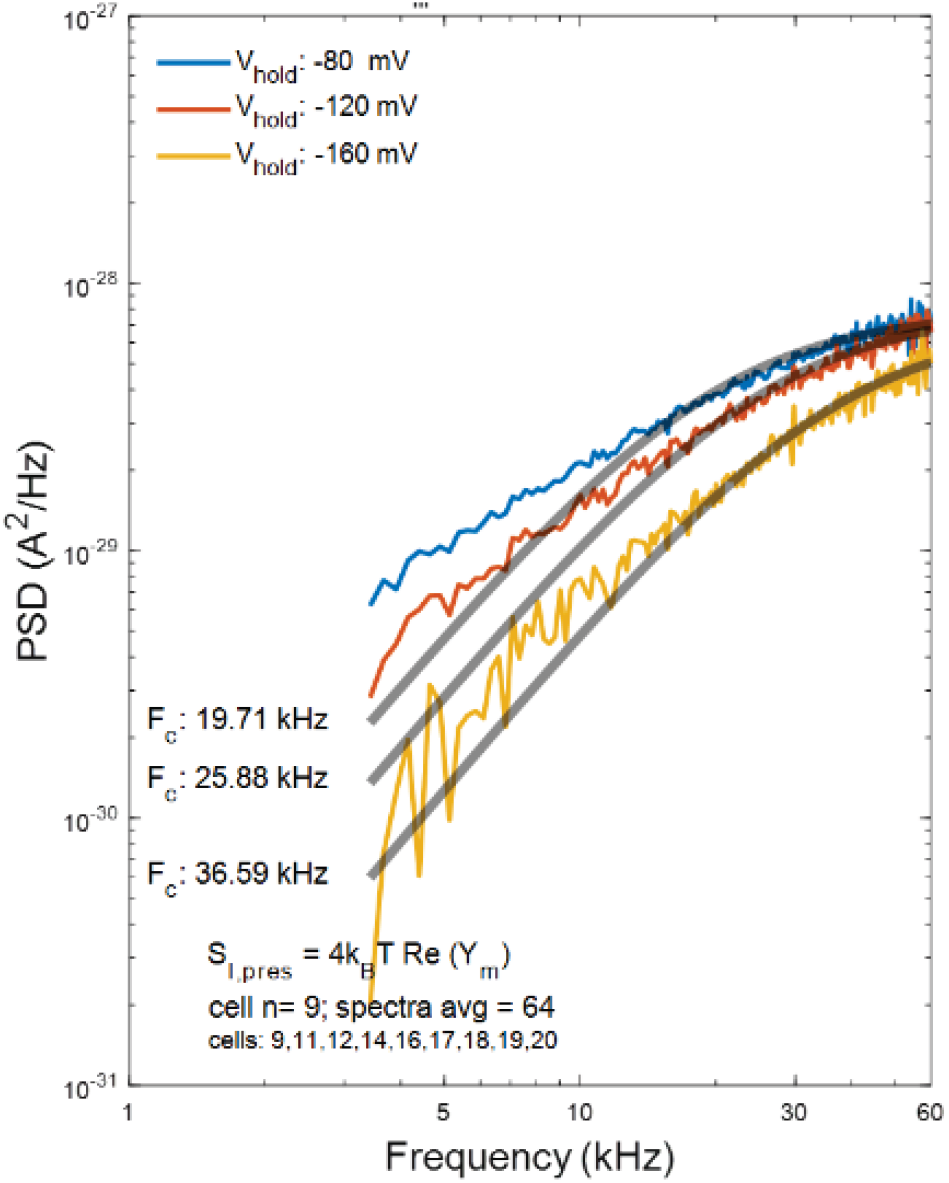
Nyquist relation noise estimates of prestin displacement currents based on membrane admittance at three holding potentials, −80 mV, −120 mV and −160 mV. Fits were made using *eq. 3* from Dong et al. (2000) (solid grey lines), which provide frequency cut-offs that increase with offset from V_h_. These cut-offs agree with those determined from **cNLC** (see **Fig. 4**).

We find that in addition to potential confounding effects of baseline headstage noise, 1/f noise invariably contributes to thermally-driven noise of OHC membrane patches (**Fig. 7**). Here, we choose 3 patches that have relatively small high frequency RF contamination below 60 kHz. In **Fig.7 A** we plot the average noise response across frequency from the three patches at holding voltages from +160 to −160 mV, showing that 1/f noise increases away from zero mV. The solid grey lines are 1/f fits to the low frequency responses. **Fig. 7 B** plots the fitted 1/f noise levels at 244 Hz as a function of voltage. Ion channel conductances are essentially absent in the OHC lateral membrane (Santos-Sacchi et al., 1997). Consequently, the main conductance in the OHC lateral membrane patch derives from prestin itself and is resistant to standard ion channel blockers (Bai et al., 2017); thus, the voltage dependent 1/f noise likely derives from prestin leakage conductance. The existence of this 1/f noise seriously interferes with extraction of the prestin noise response because it is necessary to subtract baseline noise levels at positive potentials, where prestin activity is largely absent, to extract prestin noise. Indeed, 1/f noise, headstage noise and RF noise all combine to challenge stationary assessment of physiological noise resulting from voltage-sensor activity.

**Fig. 7.**
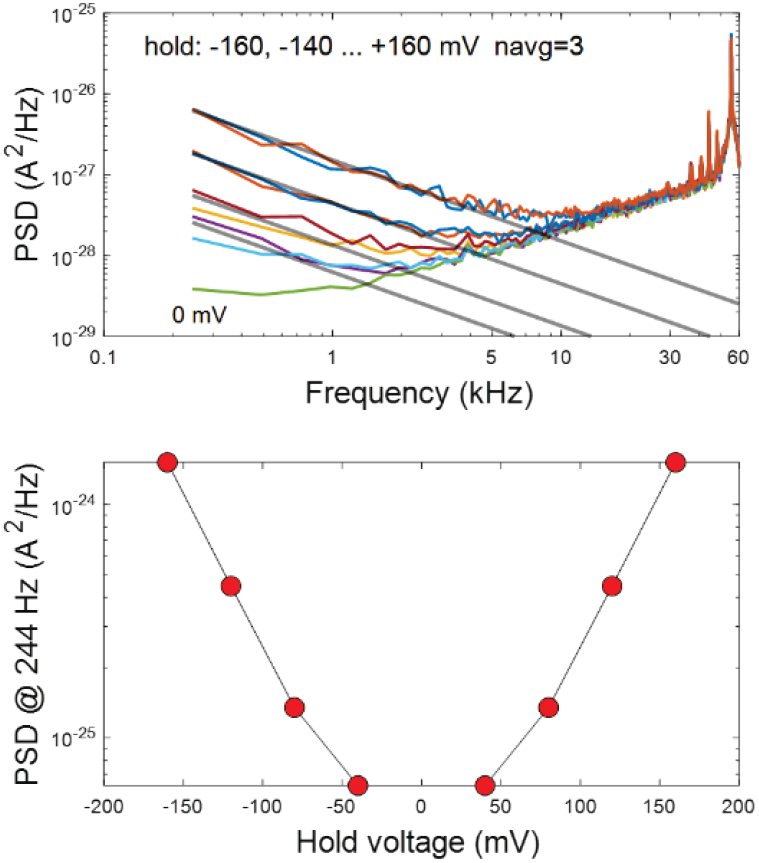
**A**) Raw stationary noise measures (colored lines) at holding potentials ranging from +160 mV to −160 mV. Some RF contamination exists at high frequencies. 1/f noise in apparent at low frequencies that is voltage-dependent and increases away from 0 mV. Grey lines are 1/f fits to the data. RF noise at high frequencies below 50 kHz (spikes) are relatively small for this group of patches. **B**) Plot of 1/f fitted response at 244 Hz, illustrating the symmetrical increase in 1/f noise away from 0 mV.

Though *non-stationary* noise analysis of ion channel gating (displacement) currents have been made and modelled [see (Catacuzzeno et al., 2021)], to our knowledge stationary noise analysis of voltage-sensor activity has not been attempted; indeed, confounding noise sources, e.g. 1/f noise, have dissuaded such attempts (Luigi Catacuzzeno, Pancho Bezanilla, personal communications). Nonetheless, Dong et al. (Dong et al., 2000) provided measures of prestin thermal noise directly (*eq. 2*) using the Axon 200B amplifier in resistive feedback mode (R_f_=0.5 GΩ), where headstage noise is quite significant. In fact, these 200B headstage noise levels are nearly twice as large as Dong et al.’s stationary noise measures across frequency (compare their data in their *Fig. 5b* with 200B headstage noise in our **Fig. 2A** or in Rosenstein’s *Fig. 3 a* (Rosenstein et al., 2012)). RF noise likely appeared in their measures, as well, since blanked high frequency regions of subtracted stationary noise data are evident in their *Fig. 5* plots. Finally, it is not clear what 1/f noise contamination they encountered, since they did not show raw, un-subtracted noise data. The frequency cut-off of their thermal noise response was several fold greater than that of their admittance-based **NLC** cut-off and expectedly, as we have shown, a corresponding Nyquist relation cut-off estimate. In our studies, we are unable to confirm their disparate observations.

**Fig. 8** illustrates raw stationary noise measures from those 3 patches in **Fig. 7** that possess different degrees of 1/f noise. 1/f noise limits the bandwidth of analyzable extracted prestin noise, since, as noted above, the spectrum at positive potentials, where **NLC** is absent, must be subtracted from spectra at other voltages. We select +120 mV spectra to subtract, rather than +160 mV spectra, to limit the untoward effect of very large 1/f contamination. Our stationary noise data (solid red and blue lines), which reduces in available bandwidth as 1/f contamination grows, cannot be reliably fit with *eq. 3* from Dong et al. (2000); however, instead we additionally overlap the corresponding Nyquist relation data (dashed red and blue lines), indicating that the two methods of noise measurement reasonably align, though inescapable RF noise interferes at the highest frequencies. No clear violation of the fluctuation-dissipation theorem appears in our measures.

**Fig. 8.**
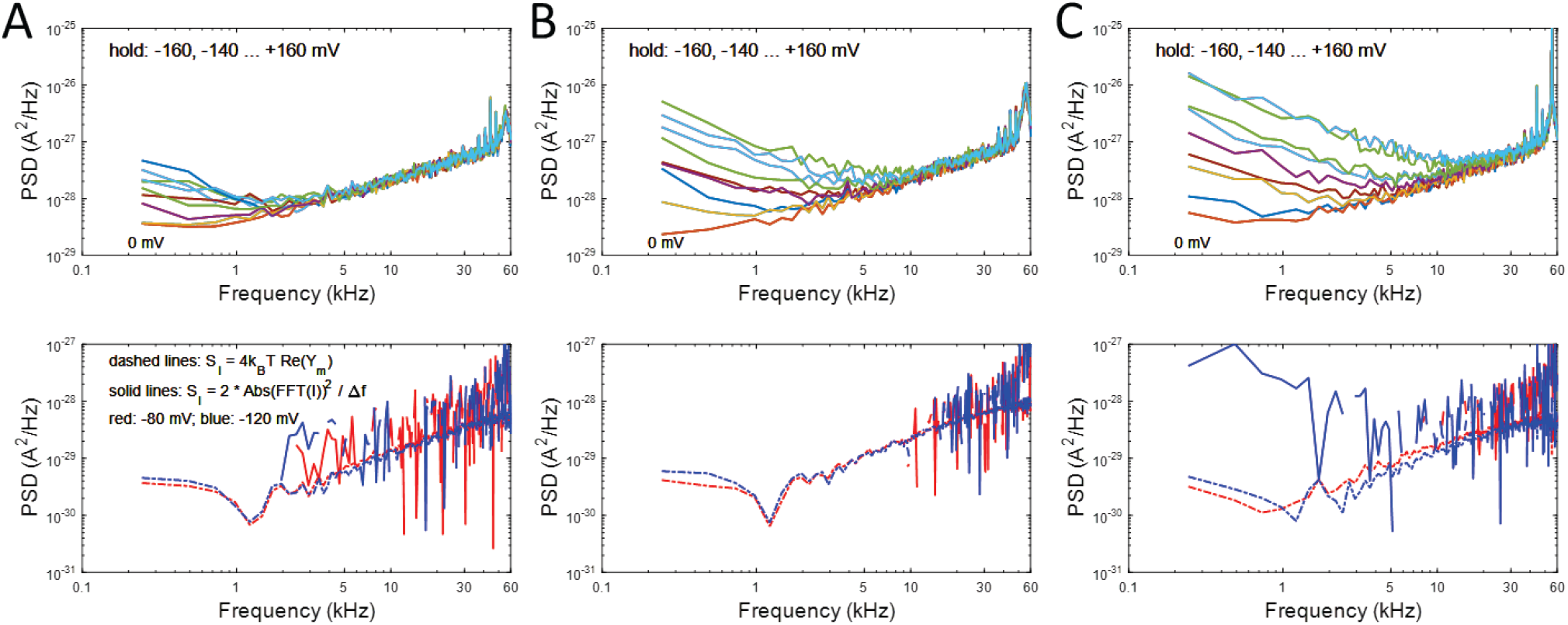
Three individual patches illustrate the difficulty extracting stationary noise in prestin. Top panels in **A, B** and **C** show raw stationary noise data collected at various holding potentials. The magnitude of 1/f noise is greater for each successive patch in **A, B** and **C**. The lower panels show prestin noise obtained by subtracting the spectrum at +120 mV from two holding voltages, −80 and −120 mV, around V_h_ (solid red and blue lines). Large 1/f noise reduces the bandwidth of available extracted prestin noise, and these data are unable to be reliably fit with *eq. 3* from Dong et al. (2000). Nevertheless, plots of the Nyquist relation noise across frequency (dashed red and blue lines) indicate that the frequency response of noise determined by either method, stationary or admittance-based, largely overlap.

To summarize, we have measured complex **NLC** out to 120 kHz, thereby enabling us to observe the rise and fall of Im(**cNLC**) across frequency, which our earlier lower bandwidth studies failed to directly reveal (Santos-Sacchi and Tan, 2022). Indeed, the high frequency fall of Im(**cNLC**) mirrors that of Abs(**cNLC**). The cut-off frequency (F_c_) of Abs(**cNLC**) corresponds to the intersection frequency (F_is_) of real and imaginary **NLC** components, confirming 2-state model expectations, see appendix in (Santos-Sacchi and Tan, 2022). Furthermore, F_is_ correlates well with displacement current noise cut-off frequency (F_c_) estimates based on the Nyquist relation (*eq. 1*), being near 19 kHz at V_h_. We were unable to determine F_c_ with fitting of directly measured stationary noise (*eq. 2*) because of 1/f noise interference. Nevertheless, our measures of voltage-driven and thermal-driven noise overlap (see **Fig. 8**). Thus, we reason that the fluctuation-dissipation theorem is not violated. Finally, voltage perturbation accurately reveals the spectral capabilities of prestin electro-mechanics, precisely because the OHC AC receptor potential, an acoustically generated voltage response, is prestin’s physiological stimulus *in vivo*.

## Acknowledgements

We thank Pancho Bezanilla and Luigi Catacuzzeno for communications concerning stationary noise analysis of voltage sensor activity in ion channels. We also thank Fred Sigworth for the use of his 200B rig to record 3 of the OHC membrane patches used in this study, the others being recorded in our own rigs. This research was supported by NIH-NIDCD R01 DC016318 and R01 DC008130.

## Notes

### Competing Interest Statement

The authors have declared no competing interest.

